# Mesolimbic opioid-dopamine interaction is disrupted in obesity but recovered by weight loss following bariatric surgery

**DOI:** 10.1101/2020.12.10.418764

**Authors:** Henry K. Karlsson, Lauri Tuominen, Semi Helin, Paulina Salminen, Pirjo Nuutila, Lauri Nummenmaa

## Abstract

**Background:** Obesity is a growing burden to health and economy worldwide. Obesity is associated with central μ-opioid receptor (MOR) downregulation, and the interaction between MOR and dopamine D_2_ receptor (D_2_R) system in the ventral striatum is disrupted among obese subjects. Weight loss recovers MOR function, but it remains unknown whether it also recovers aberrant opioid-dopamine interaction. Here we addressed this issue by studying subjects undergoing surgical weight loss (bariatric surgery) procedure.

**Methods:** We recruited 20 healthy non-obese (mean BMI 22) and 25 morbidly obese women (mean BMI 41) eligible for bariatric surgery. Brain MOR and D_2_R availability was measured using positron emission tomography (PET) with [^11^C]carfentanil and [^11^C]raclopride, respectively. Either Roux-en-Y gastric bypass or sleeve gastrectomy was performed to obese subjects according to standard clinical treatment. 21 obese subjects participated in the postoperative PET scanning six months after bariatric surgery.

**Results:** In the control subjects, MOR and D_2_R availabilities were associated in the ventral striatum (*r* = .62) and dorsal caudate (*r* = .61). Preoperatively, the obese subjects had disrupted association in the ventral striatum (*r* = .12) but unaltered association in dorsal caudate (*r* = .43). The association between MOR and D_2_R availabilities in the ventral striatum was recovered (*r* = .62) among obese subjects following the surgery-induced weight loss (mean total weight loss 22 %).

**Conclusions:** Bariatric surgery and concomitant weight loss recovers the interaction between MOR and D_2_R in the ventral striatum in the morbidly obese. Consequently, the dysfunctional opioid-dopamine interaction in the ventral striatum is likely associated with an obese phenotype and may mediate excessive energy uptake. Striatal opioid-dopamine interaction provides a feasible target for pharmacological and behavioural interventions for treating obesity.

**Clinical Trials Registration:** SleevePET2, NCT01373892, http://www.clinicaltrials.gov

## Introduction

The prevalence of obesity is dramatically increasing and there is urgent need for novel efficient therapies. Numerous studies point towards the role of brain in the development and maintenance of obesity (Stice and Burger, 2019; Morales and Berridge, 2020). Previous studies indicate that both opioid and dopamine systems in brain’s reward circuit are dysfunctional in obesity. Endogenous opioid system has been linked to hedonic aspects of feeding in animals (Pecina and Smith, 2010; Fields and Margolis, 2015). In humans, both μ-opioid receptor (MOR) antagonists and inverse agonists have been shown to reduce eating behaviour (Nathan et al., 2012; Cambridge et al., 2013). Previously, decreased MOR availability has been observed in reward circuit among obese subjects (Burghardt et al., 2015; Karlsson et al., 2015). Thus, aberrant opioid functioning in obesity may diminish the opioid dependent rewarding effects of eating. Alterations in dopamine D_2_ receptor (D_2_R) expression and function in obesity has been observed in some (Volkow et al., 2008; Stice et al., 2011; Salamone and Correa, 2013; Lee et al., 2018) but not all human imaging studies (Eisenstein et al., 2013; Karlsson et al., 2015). It is possible that the relationship between D_2_R availability and measures of obesity is not linear, but quadratic (Horstmann et al., 2015) or age-dependent (Dang et al., 2016). Alternatively, it is possible that the effects of obesity on D2R are mediated via the MOR system.

Tight interaction between dopaminergic and opioidergic systems has been proposed to underlie human reward functions (Le Merrer et al., 2009), but only a few studies have actually investigated this issue. In humans, dopamine-releasing drugs such as cocaine and amphetamine lead to endogenous opioid release (Soderman and Unterwald, 2009; Colasanti et al., 2012; Mick et al., 2014). In rats, both D_2_Rs and MORs are closely connected in the striatum, which can be morphologically divided into striosome/patch and matrix compartments. MORs can control the release of dopamine through inhibiting GABAergic interneurons in VTA (Spanagel et al., 1992; Kalivas, 1993; Volkow and Wise, 2005; Rada et al., 2010; Jalabert et al., 2011; Chartoff and Connery, 2014). Furthermore, VTA dopamine neurons express MOR postsynaptically, and direct inhibition between MOR and dopamine neurons exists without GABAergic signaling (Margolis and Hjelmstad, 2014).

Cross-talk between opioidergic and dopaminergic system may underlie aberrant reward-related behaviours, such as excessive feeding. In rats, intravenous administration of MOR agonists triggers dopamine release and feeding (Yeomans and Gray, 2002), while MOR antagonists block dopamine release and reduce food consumption (Taber et al., 1998). Finally, *in vivo* PET data from humans show that there is a close interaction between MOR and D_2_R receptors in the reward circuit among non-obese subjects, while this interaction is disrupted in the ventral striatum among obese subjects, potentially contributing to obesity (Tuominen et al., 2015). However, it remains unclear whether the dysfunctional MOR / D_2_R interaction reflects a vulnerability endophenotype for obesity, or whether it develops as a consequence of the obese state.

Bariatric surgery is the most effective method for weight loss in obesity (Adams et al., 2017). The surgical procedure significantly lowers appetite (Bryant et al., 2020), but the actual molecular brain mechanisms behind this are still poorly understood. Bariatric surgery provides a powerful method for investigating changes in neuroreceptor systems and opioid-dopamine interaction after weight gain. Previous studies have investigated effects of bariatric surgery and following weight loss to separate receptor systems, showing mainly unaltered D_2_R availability and normalized MOR availability (Dunn et al., 2010; Steele et al., 2010; de Weijer et al., 2014; Karlsson et al., 2016). Here we tested whether bariatric surgery induced weight loss could recover the dysfunctional opioid-dopamine interaction in the obese.

## Materials and Methods

The study was conducted in accordance with the Declaration of Helsinki and approved by the Ethical Committee of the Hospital District of South-Western Finland (SleevePET2, NCT01373892, http://www.clinicaltrials.gov). All participants gave signed informed consent form prior to scans.

### Subjects

We recruited 25 morbidly obese women (mean BMI 41 kg/m^2^) eligible for bariatric surgery. Either Roux-en-Y gastric bypass or sleeve gastrectomy was performed as their standard clinical treatment. Four subjects discontinued the study for personal reasons, and 21 participated in the postoperative scanning six months after the surgical procedure. 20 non-obese healthy women (mean BMI 22 kg/m^2^) formed the control group. Data for this patient cohort have been reported previously (Karlsson et al., 2015; Tuominen et al., 2015; Karlsson et al., 2016). Characteristics of the subjects are presented in the Table 1. Clinical screening of the subjects included history, physical examination, anthropometric measurements, and laboratory tests. Exclusion criteria involved opiate drug use, binge-eating disorders, neurological and severe mental disorders, substance abuse, excessive alcohol consumption (more than eight units per weeks) determined by clinical interviews, medical history, and blood tests. None of the controls smoked tobacco, but 8 obese subjects were smokers (3-15 cigarettes per day). Antidiabetic, antihypertensive, and cholesterol lowering drugs were paused prior to the study.

**Table 1.** Characteristics of the subjects. Data are presented as mean ± SD.

|  | Obese preoperative (N = 25) | Obese postoperative (N = 21) | Healthy control subjects (N = 20) |
| --- | --- | --- | --- |
| Age (y) | 41.2 ± 9.2 | - | 42.0 ± 13.2 |
| BMI (kg/m<sup>2</sup>) | 41.3 ± 4.1 | 31.9 ± 4.4 | 22.4 ± 2.6 |
| Percentage of fat (%) | 50.3 ± 6.7 | 43.2 ± 4.2 | 30.6 ± 6.4 |
| Tobacco smokers / non-smokers (N) | 8 / 17 | 5 / 16 | 0 / 20 |
| Amount of alcohol use (units per week) | $1.7 \pm 1.8$ | N / A | $2.9 \pm 2.3$ |
| Injected activity of [$^{11}\text{C}$]carfentanil (MBq) | $253.2 \pm 11.6$ | $252.1 \pm 15.0$ | $251.2 \pm 8.4$ |
| Injected activity of [$^{11}\text{C}$]raclopride (MBq) | $247.9 \pm 20.8$ | $254.5 \pm 10.9$ | $258.3 \pm 15.7$ |

### Image Acquisition and Quantification of Receptor Availability

We measured D_2_ receptor availability with the antagonist [^11^C]raclopride (Farde et al., 1986) and μ-opioid receptor availability with the high-affinity agonist [^11^C]carfentanil (Frost et al., 1985) using positron emission tomography (PET) on two separate visits. Subjects were scanned again with both radiotracers six months after bariatric surgery. Radiotracer production has been described previously (Karlsson et al., 2015). Both radioligands had high radiochemical purity (>99 %). Before scanning, a catheter was placed in the subject’s left antecubital vein for tracer administration. Head was strapped to the scanner table in order to prevent head movement. Subjects fasted two hours prior to scanning. A CT scan was performed to serve as attenuation map. Clinical well-being of subjects were monitored during the scanning.

We injected both tracers as bolus in separate scans on separate days. Injected amounts of [^11^C]carfentanil and [^11^C]raclopride are presented in the Table 1. After injection, radioactivity in brain was measured with the GE Heatlhcare Discovery™ 690 PET/CT scanner (General Electric Medical Systems, Milwaukee, WI, USA) for 51 minutes, using 13-time frames. MR imaging was performed with Philips Gyroscan Intera 1.5 T CV Nova Dual scanner to exclude structural abnormalities and to provide anatomical reference images for the PET scans. Anatomical images (1 mm^3^ voxel size) were acquired using a T1-weighted sequence (TR 25 ms, TE 4.6 ms, flip angle 30°, scan time 376 s).

All alignment and coregistration steps were performed using SPM8 software (www.fil.ion.ucl.ac.uk/spm/) running on Matlab R2012a (The Mathworks Inc., Sherborn, Massachusetts). To correct for head motion, dynamic PET images were first realigned frame-to-frame. The individual T1-weighted MR images were coregistered to the summation images calculated from the realigned frames. Regions of interest (ROIs) for reference regions were drawn manually on MRI images using PMOD 3.4 software (PMOD Technologies Ltd., Zurich, Switzerland). Occipital cortex was used as the reference region for [^11^C]carfentanil and cerebellum for [^11^C]raclopride. Receptor availability was expressed in terms of *BP*_ND_, which is the ratio of specific to non-displaceable binding in brain. *BP*_ND_ was calculated applying basis function method for each voxel using the simplified reference tissue model (SRTM) with reference tissue time activity curves (TAC) as input data (Gunn et al., 1997).

The subject-wise parametric *BP*_ND_ images were normalized to the MNI space using the T1-weighted MR images, and smoothed with a Gaussian kernel of 8 mm FWHM. Anatomic regions of interest were generated in the ventral striatum, dorsal caudate nucleus, and putamen using the AAL (Tzourio-Mazoyer et al., 2002) and Anatomy (Eickhoff et al., 2005) toolboxes. Statistical analysis was performed as described earlier (Tuominen et al., 2015). In the ROI analysis, Pearson correlation was calculated between the tracer-wise *BP*_ND_s in the striatal regions of interest. Fisher’s z-test was used for quantifying whether ROI-level Pearson correlations between the [^11^C]raclopride and [^11^C]carfentanil *BP*_ND_ values were statistically different between groups.

## Results

MOR and D_2_R availabilities were associated in the ventral striatum (*r* = .62, p < 0.05) and dorsal caudate (*r* = .61, p < 0.05) in the control subjects (Figure 1). Preoperatively, the obese subjects had disrupted association in the ventral striatum (*r* = .12, ns), but unaltered association in dorsal caudate (*r* = .43, p < 0.05) (Figure 1). MOR and D_2_R availabilities in putamen were not associated in either group.

**Figure 1.**
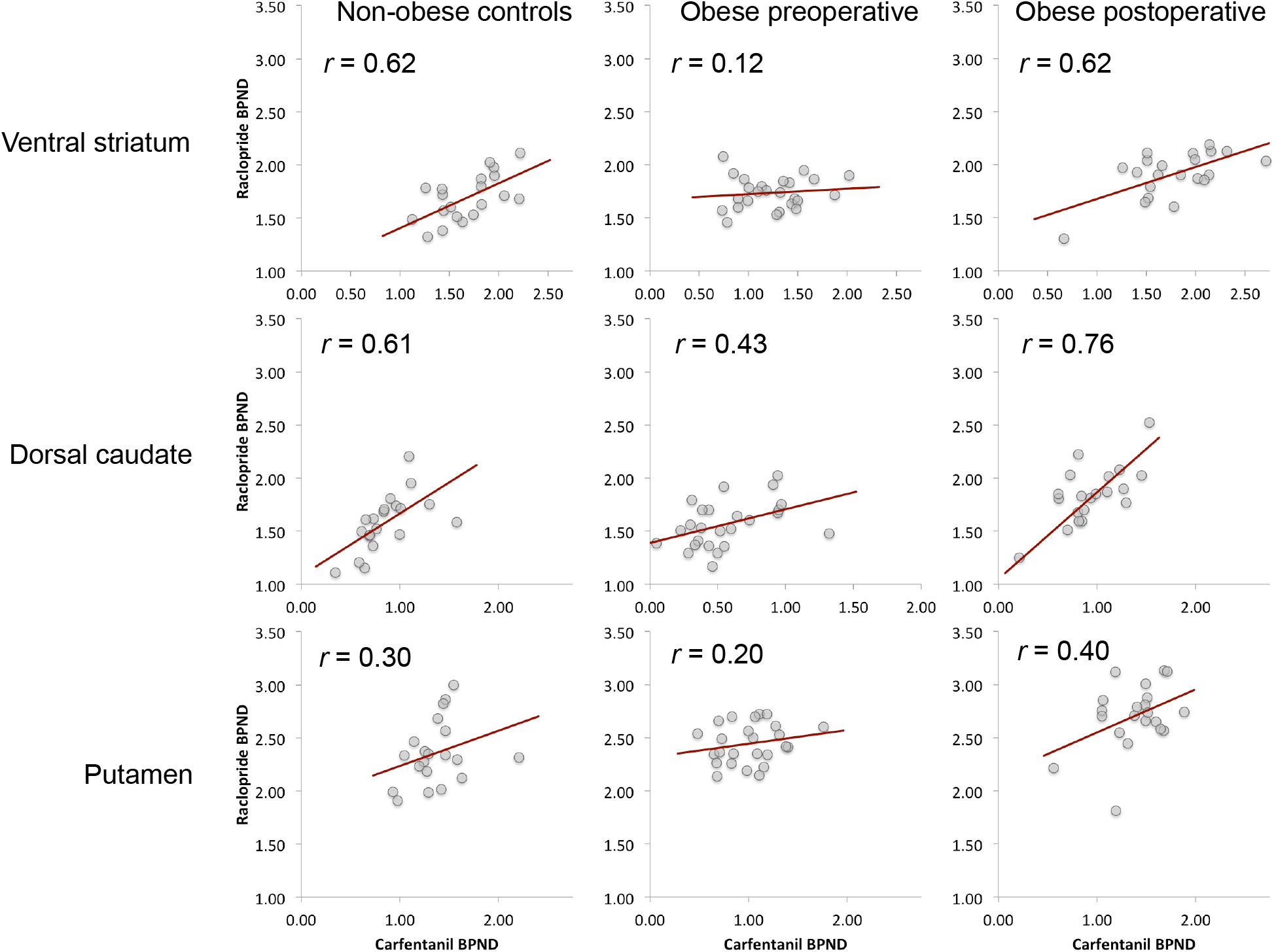
Correlations between [^11^C]raclopride *BP*_ND_ and [^11^C]carfentanil *BP*_ND_ in the ventral striatum, dorsal caudate, and putamen. The association is recovered in the ventral striatum after surgery-induced weight loss. Significant association was found in dorsal caudate in all groups, whereas no association was found in putamen in any group.

The association between MOR and D_2_R availabilities in the ventral striatum was recovered (*r* = .62, p < 0.05) among obese subjects following the surgery-induced weight loss (mean total weight loss 25.0 ± 8.2 kg and 22.1 ± 6.1 %) (Figure 1 and 2). There was no difference between the two surgical procedures in receptor availabilities or the association between receptors before or after surgery.

**Figure 2.**
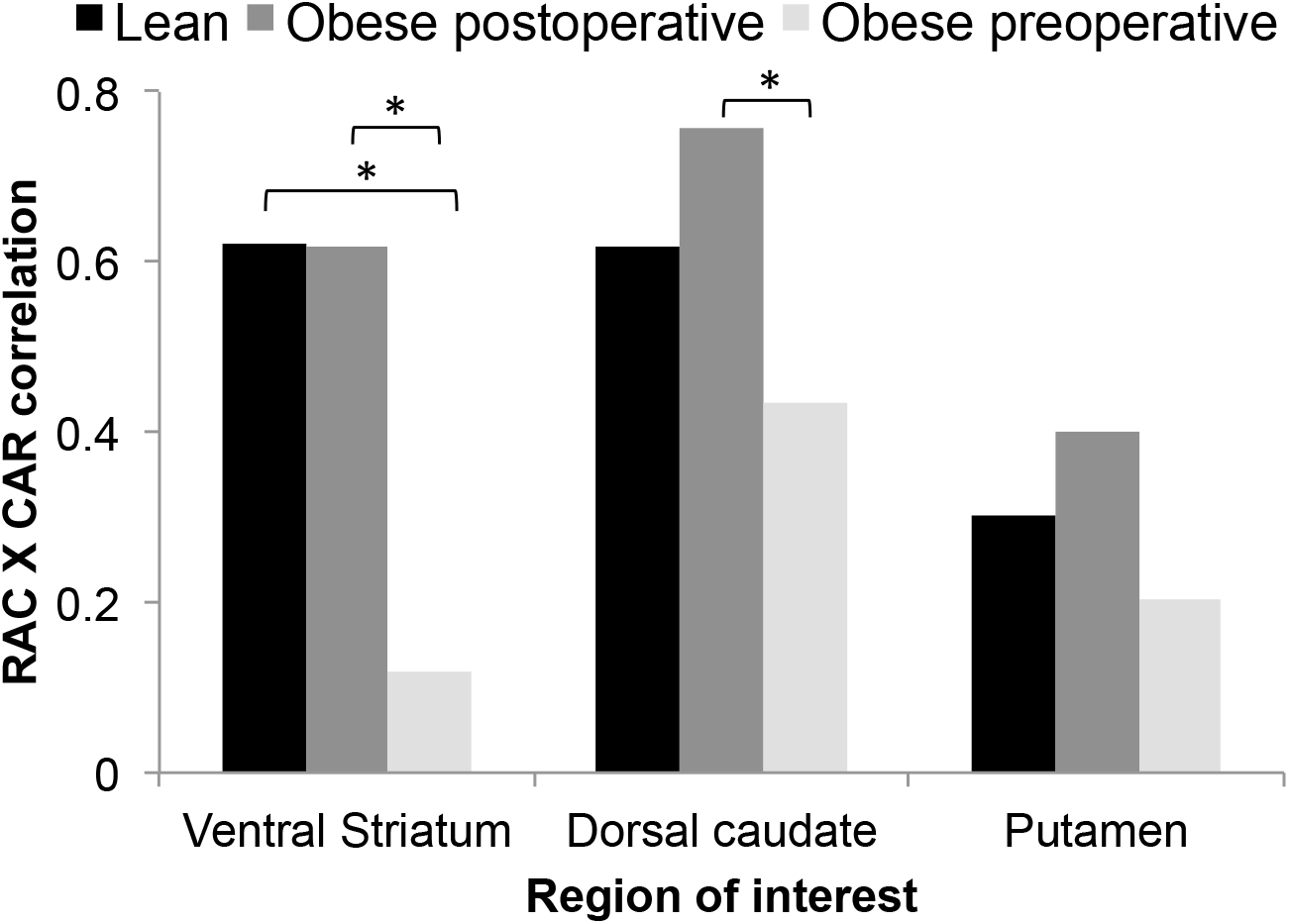
Mean correlations between [^11^C]raclopride *BP*_ND_ and [^11^C]carfentanil *BP*_ND_ in non-obese and obese subjects before and after surgery. Asterisks denote significant between-groups differences.

## Discussion

Our main finding was that opioid-dopamine interaction is recovered by bariatric surgery and concomitant weight loss. Dysfunctional opioid-dopamine interaction in the ventral striatum is associated with an obese phenotype and may mediate excessive energy uptake, and we have reported earlier that MOR levels return to normal after weight loss (Karlsson et al., 2016). Behaviorally this is in line with previous studies, showing improved satiety and lowered appetite after bariatric surgery (Morinigo et al., 2006; Karamanakos et al., 2008). We have previously shown that striatal opioid and dopamine systems are coupled in non-obese but not in obese subjects (Tuominen et al., 2015). In the normal-weight subjects the interaction was strongest in the ventral striatum, but also significant in dorsal caudate. Growing evidence indicates that MOR and D_2_R are expressed in same striatal neurons (Ambrose et al., 2004; Margolis and Hjelmstad, 2014). The interaction between these receptor systems is likely crucial in regulating appetite, because it breaks down in the striatum in the obese subjects, while association in the dorsal caudate remains intact. This might explain unaltered D_2_R levels in obesity: although obesity-dependent dysfunction in dopaminergic system is shown in numerous animal studies, it may be mediated through MOR-dependent mechanisms without having any effect on the actual number of D_2_R proteins. Even if the amount of D_2_R protein stays the same in obesity, decoupling of MOR and D_2_R in the striatum may cause altered dopaminergic functions.

MORs are co-localized with D_2_Rs in striosomes (Herkenham and Pert, 1981). Dopaminergic neurons in the striosomes project directly to ventral tegmental area (VTA) and substantia nigra, whereas neurons projecting to GABAergic neurons are distributed in the matrix compartment (Watabe-Uchida et al., 2012). Pathways projecting from striosomes back to the midbrain exert disinhibitory control over the dopaminergic neurons (Fujiyama et al., 2015), thus having direct influence on the reward functions. These neurons are under direct opioidergic control (Banghart et al., 2015). Accordingly, endogenous opioids disinhibit the neurons projecting from the patches to the midbrain (i.e. disinhibiting the disinhibiting neurons), and in this way increase dopaminergic firing in VTA. The rewarding effects of opioids are dependent on the MORs located in the striosomes (Cui et al., 2014)

Based on the observation that dopamine release caused by opioids in the striatum is dependent on the MORs in the striosomes in mice (Cui et al., 2014), we hypothesize that aberrant opioid function in obese humans might lead to diminished dopamine release caused by eating. When obese subjects lose weight, the interaction between MOR and D_2_R is reverted. This further supports the notion that the interaction between these receptor systems is a normal state. The dysfunction of opioid-dopamine interaction in the ventral striatum might be an important factor underlying overeating, and thus a feasible target for pharmacological and behavioural interventions. This has already been noted in pharmacological studies. MOR antagonist naltrexone therapy alone does not lead to significant weight loss, but promising results are obtained when it is coupled with bupropion (a dopamine and norepinephrine re-uptake inhibitor) (Lee and Fujioka, 2009; Greenway et al., 2010; Smith et al., 2013; Billes et al., 2014). Combination therapy of naltrexone and bupropion has been approved by FDA and EMA for weight management in adults (Sherman et al., 2016) and certain amount of obese patients achieve significant weight loss (Fujioka et al., 2016; Mullally et al., 2020; Onakpoya et al., 2020). The favourable effect of the combination therapy may be due to tight coupling of MOR and D_2_R. Moreover, the better efficacy of the combination therapy over monotherapies underlines the complex pattern of neurotransmitter networks underlying overeating, and suggests that both aspects of reward functions – *wanting* and *liking*, processes mediated by dopaminergic and opioidergic systems, respectively (Berridge and Kringelbach, 2015) – have to be taken care of in order to treat obese patients.

### Limitations of the study

Only female subjects were studied, and the results may not be generalizable to male subjects. It was not possible to differentiate the combined effects of postoperative weight loss and altered gut anatomy and function. Altered neuroreceptor interaction may be due to the changes in gut hormones but also due to reduced intake of palatable foods. Further studies are needed to elucidate the sole effect of weight loss due to altered energy intake on interaction of opioid and dopamine receptors by comparing the effects of weight loss by surgery versus dieting.

### Conclusions

Obesity is associated with disrupted opioid-dopamine interaction in the ventral striatum, but this is recovered by weight loss after bariatric surgery. The dysfunction of opioid-dopamine interaction might be an important factor underlying overeating.

## Acknowledgments

The study was conducted within the Finnish Centre of Excellence in Cardiovascular and Metabolic Diseases supported by the Academy of Finland (grants #251125, #121031, #304385), Sigrid Juselius Foundation, University of Turku, Turku University Hospital and Åbo Akademi University. HKK was supported by personal grants from The Finnish Diabetes Research Foundation and The National Graduate School of Clinical Investigation. The funders had no role in study design, data collection and analysis, decision to publish, or preparation of the manuscript. The authors thank the staff of the Turku PET Centre for assistance in the PET imaging. Special thanks goes to research nurse Mia Koutu.

